# MoMI-G: Modular Multi-scale Integrated Genome Graph Browser

**DOI:** 10.1101/540120

**Authors:** Toshiyuki T. Yokoyama, Yoshitaka Sakamoto, Masahide Seki, Yutaka Suzuki, Masahiro Kasahara

## Abstract

Genome graph is an emerging approach for representing structural variants on genomes with branches. For example, representing structural variants of cancer genomes as a genome graph is more natural than representing such genomes as differences from the reference genome. However, there is currently no visualization method for large genome graphs, such as human cancer genomes. To this end, we developed MOdular Multi-scale Integrated Genome graph browser, MoMI-G, a web-based genome graph browser that can visualize genome graphs with structural variants and supporting evidences such as read alignments, read depth, and annotations.

## INTRODUCTION

Structural Variants (SVs), which are often characterized as 50 bp or larger genomic rearrangements of chromosomal segments, are associated with various human diseases [1–3]. For example, some fusion genes caused by SVs are known oncogenes [4]. Identifying SVs and interpreting their potential impacts are critical steps toward cataloguing the variations in the human genome and mechanistic understanding of genetic diseases and cancers.

SV visualization is a very important step in an SV calling process, because it enables the manual inspection of SVs for achieving two goals: The first is to better understand the relationships between SVs and other genomic features, and the second is to ensure a smaller number of false positives.

For the first goal, SV visualization tools should be able to simultaneously display multiple intervals along with their relationships, even when the breakpoints are distant or when SVs are nested. Older SV visualization tools focus on visualizing only canonical SVs (insertion, deletion, inversion, duplication, and translocation) [5,6], because they account for a significant portion of the identified SVs at that time. However, as long-read sequencing technologies reveal an increasing number of SVs, SV visualization with the existing tools becomes more challenging. For example, a large inversion is often identified as two separate translocations at the two breakpoints of the inversion; one might not be able to immediately recognize that the two translocation events are explained by a single large inversion. Another example is a nested SV. When there is a large inversion that contains several smaller SVs, such as insertion of transposons or deletions, the nested SVs often obscure the relationship between genomic regions that are distant in the reference genome, but are actually close in the target genome. To this end, we employed genome graphs as a theoretical backbone for providing more systematic way of presenting SVs with varying complexities, including nested and large SVs [7]. Genome graphs can represent SVs more naturally than those that represent SVs as differences from a reference genome (e.g., VCF). However, there is no visualization method for large genome graphs, such as human cancer genomes.

For the second goal, manual inspection of SVs identified using SV calling tools is important, because these tools are not yet accurate enough; human experts are required to accurately and reliably distinguish true positive SVs from false positive ones. For example, SVs identified by using reads obtained by different sequencing platforms are often not concordant [8], suggesting that the algorithms of SV callers need further improvement. Therefore, SV candidates need to be manually inspected using read alignments and genomic annotations [9]. Developers of SV callers may wish to track down false positives, so they can improve the algorithms. Biologists may wish to filter out false positives through manual inspection to find genuine causal SVs. However, manual inspection by using existing SV visualization tools occasionally becomes very difficult for certain cases. For example, for nested SVs and long reads spanning over multiple breakpoints, existing tools cannot show the read alignments in multiple intervals at a glance, making it unrealistic to manually judge the authenticity of candidate SVs. Another example is that tandem duplications identified by SV callers are often inaccurate. This is presumably because the accuracy for finding tandem duplications has not been optimized for real biological data [8,10,11].

To achieve these two goals, we developed MoMI-G (pronounced mo-me-jee), a genome graph browser that visualizes SVs using variation graphs (Fig. 1, Additional file 1: Supplemental Fig. 1), a variant of genome graphs [12]. Herein, we describe the use cases and features of MoMI-G using the LC-2/ad human lung adenocarcinoma cell line that carries a CCDC6-RET fusion gene [13–16], and CHM1, a human hydatidiform mole cell line that originates from a single haploid [17]. MoMI-G helps in understanding the entire picture of SVs, even those that are nested or large, regardless of their size. MoMI-G allows researchers to obtain novel biological knowledge by comparing a reference genome with an individual genome by using a variation graph.

**Figure 1.**
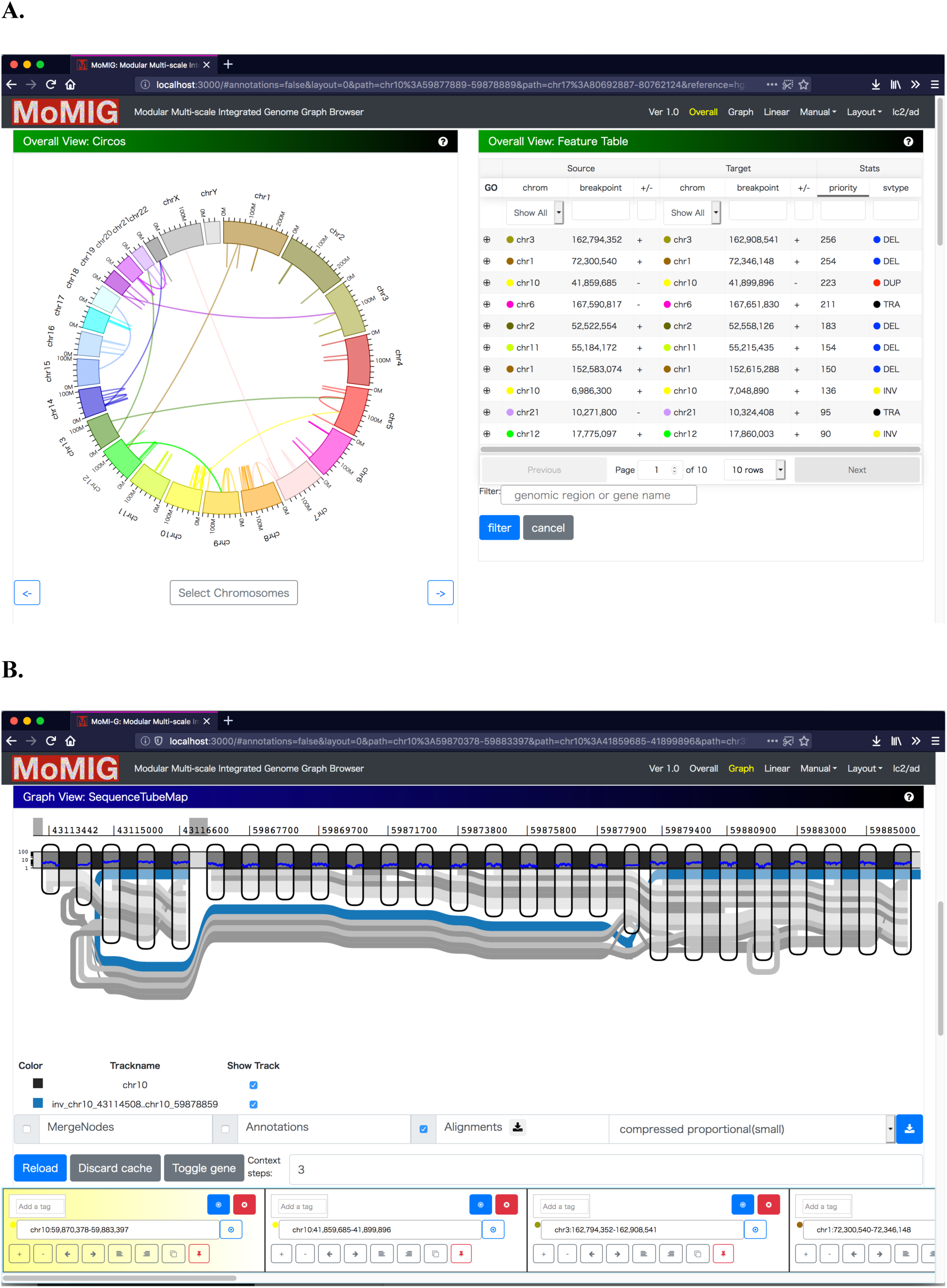
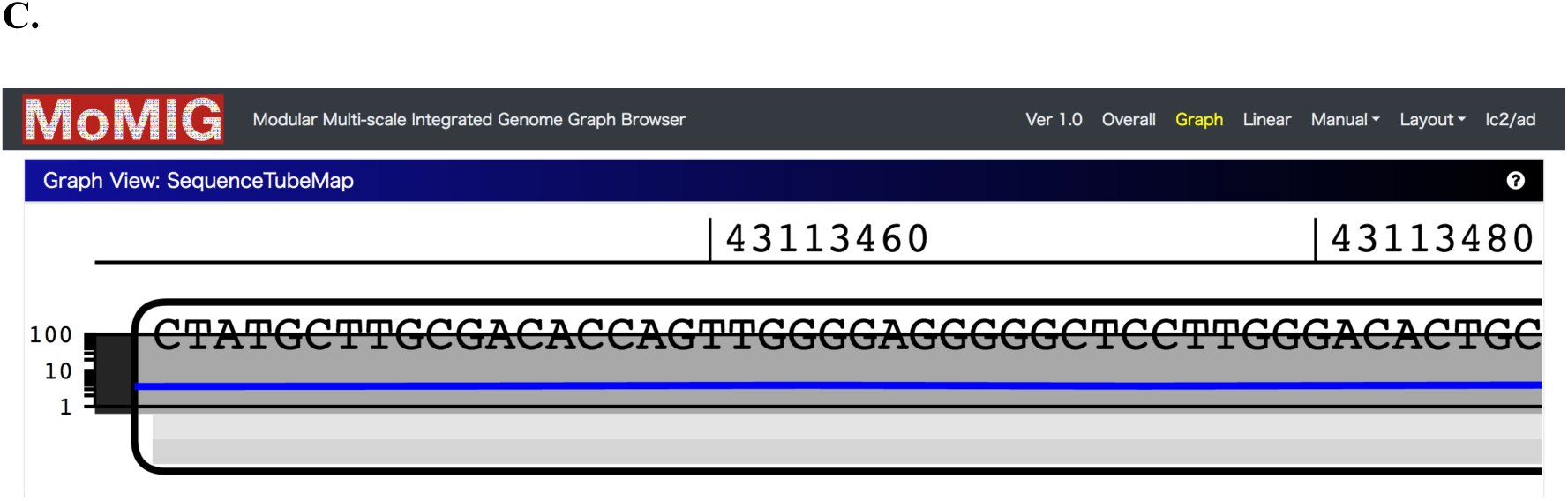
Overview of MoMI-G. A user typically selects one of the preset combinations of view modules. The user can customize the window by adding or removing view modules, if necessary. Three examples showing views of different scales are shown. Comprehensive descriptions for all modules are shown in Additional file 1: Supplemental Fig. 1. (A) Chromosome-scale view: Circos Plot (left) shows the distribution of SVs over all chromosomes. Arcs are chromosomes. Curves represent SVs. Feature Table (right) shows a filtered/sorted list of the SVs in an input VCF file. (B) Gene-scale view: SequenceTubeMap (top) shows the graphical view of the genomic region selected in Circos Plot, Feature Table, or Interval Card Deck. A rounded rectangle is a node that represents a piece of a genomic sequence. The thick lines spanning over nodes are paths; the horizontal thick black line with light/dark shades is a chromosome of the reference genome, and the blue line indicates one end of an inversion. The color of lines indicating SVs is assigned arbitrarily. Read alignments are shown as gray thin lines, suggesting that the inversion here is likely heterozygous. Interval Card Deck (bottom) queues a list of genomic intervals for candidate SVs selected by using Circos Plot or Feature Table for rapidly screening hundreds of candidate intervals. (C) Nucleotide-scale view: SequenceTubeMap can show nucleotides.

## RESULTS

MoMI-G is a web-based genome browser developed as a single-page application implemented in TypeScript and with React. Because users need different types of views, even for the same data, MoMI-G provides three groups of view modules for the analysis of SVs at different scales, namely chromosome-scale, gene-scale, and nucleotide-scale view groups (Additional file 1: Supplemental Table 1). Users can use one or more view modules in a single window.

The input of MoMI-G is a variation graph, read alignment (optional), and annotations (optional). MoMI-G accepts a succinct representation of a vg variation graph, which is an XG file, as a variation graph. A script that converts a FASTA file of a reference genome and a common variant format (VCF) file into an XG file is included in the MoMI-G package, although the VCF format cannot represent some types of SVs that the XG format can represent, such as nested insertions. Read alignment data on the graph need to be represented as a graph alignment map (GAM) file; alternatively, users can convert a binary alignment map (BAM) file into a GAM file, although this is not recommended due to some limitations.

We show three examples from two samples to demonstrate the utility of MoMI-G. One of the examples is a large inversion, and the other is nested SVs that are difficult to visualize using existing tools. We also show nested SVs from the CHM1 dataset [17]. For all the examples, we used the Amazon EC2 instance type t2.large with 8 GB of memory and 2.4 GHz Intel Xeon processor as the MoMI-G server; the server requirement for this tool is minimal. MoMI-G supports common browsers, including Chrome, Safari, and Firefox.

### Data model used in MoMI-G and MoMI-G tools

To our knowledge, no publicly available SV visualization tools are available for large and nested SVs with alignments of long reads. Thus, we aimed to develop a genome graph browser at the earliest so that users can obtain new biological knowledge from real data. We used an existing library, SequenceTubeMap (https://github.com/vgteam/sequenceTubeMap), for visualizing a variation subgraph, rather than developing our own library from scratch.

SequenceTubeMap is a JavaScript library that visualizes multiple related sequences such as haplotypes. A variation graph used in SequenceTubeMap is a set of nodes and paths, where a node represents part of a DNA sequence, and a path represents (part of) a haplotype. Edges are implicitly represented by adjacent nodes in paths.

MoMI-G accepts variation graphs in which SVs are represented by paths so that SequenceTubeMap can visualize them. A deletion is represented by a path that skips over a sequence that other paths pass through. Similarly, an insertion is represented by a path that passes through an extra sequence that other paths do not visit; an inversion is represented by a path where part of the sequence in other paths is reversed; and, a duplication is represented by a path that passes through the same sequence twice or more.

The MoMI-G package includes a set of scripts (MoMI-G tools) that convert a VCF file into the variation graph format. We used MoMI-G tools for generating the input variation graphs; alternatively, users can generate variation graphs on their own. See the method section and Additional file 1: Supplemental Figure 2 for details of the MoMI-G tools. Briefly, MoMI-G tools convert a VCF record into a path in the output variation graph. A deletion is converted into a path that starts, at most, 1 Mbp before one breakend of the deletion, traverses to the breakend, jumps to the other breakend, and proceeds for a certain length (<1 Mbp). Note that the sequences flanking the deletion are added to indicate the edge representing the deletion because edges are implicitly represented in SequenceTubeMap. Insertions, inversions, and duplications are similarly represented by paths with flanking sequences.

### Visualization Examples

#### Revealing a large SV: A large inversion and a subsequent short deletion

Using MoMI-G, we show an example of a complex SV that involves two SVs identified by Sniffles, each of which connects two different points on a chromosome. This complex SV can be considered a large inversion and two flanking deletions. Previous studies involving the use of whole genome sequencing or RNA-seq with the Illumina HiSeq or Nanopore MinION identified the CCDC6-RET fusion gene in LC-2/ad [14–16,18]. However, those studies focused only on the region around the CCDC6-RET fusion point, and the entire picture, including the other end of the inversion, was unclear. To address this issue, we explored the wider region around CCDC6-RET with MoMI-G.

First, we sequenced the genome of LC-2/ad with Oxford Nanopore MinION R9.5 pore chemistry and merged reads with those from a previous study (accession No. DRX143541-DRX143544) [18]. We generated 3.5 M reads to 12.8× coverage in total and then aligned them with GRCh38. The average length of the aligned reads was 16 kb (Additional file 1: Supplemental Table 2). We detected 11,316 SVs in the VCF format, including the previously known CCDC6-RET fusion gene, on the nuclear DNA of LC-2/ad cell line (Additional file 1: Supplemental Table 3). See the methods section for details.

The distance between RET (chr10: 43,075,069−43,132,349) and CCDC6 (chr10: 59,786,748−59,908,656) is about 17 Mbp in GRCh38. We confirmed that a CCDC6-RET fusion gene exists in LC-2/ad (Fig. 2A). This fusion gene is presumably caused by an inversion, although only one end of the inversion was found. We found an unknown novel adjacency that well explains the other end of the inversion (Fig. 2B, Additional file 1: Supplemental Table 4). MoMI-G was able to display the relationships between the two breakends of the inversion, enabling users to understand large SVs. We explored the read alignments around the fusion and found that the fusion was heterozygous (Fig. 2C). MoMI-G is the first stand-alone genome graph browser that can display long-read alignments over branching sequences that represent a heterozygous SV.

**Figure 2.**
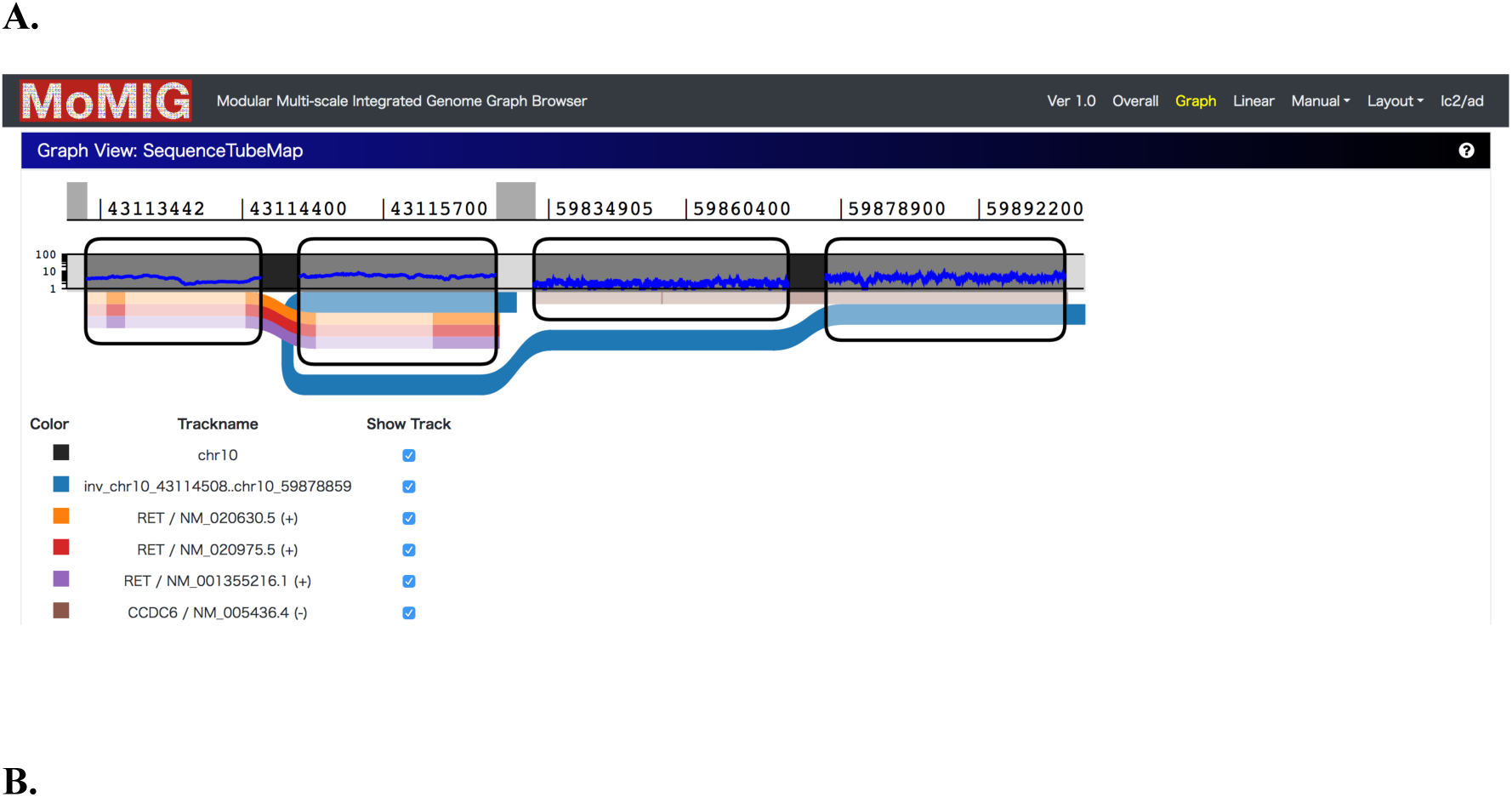
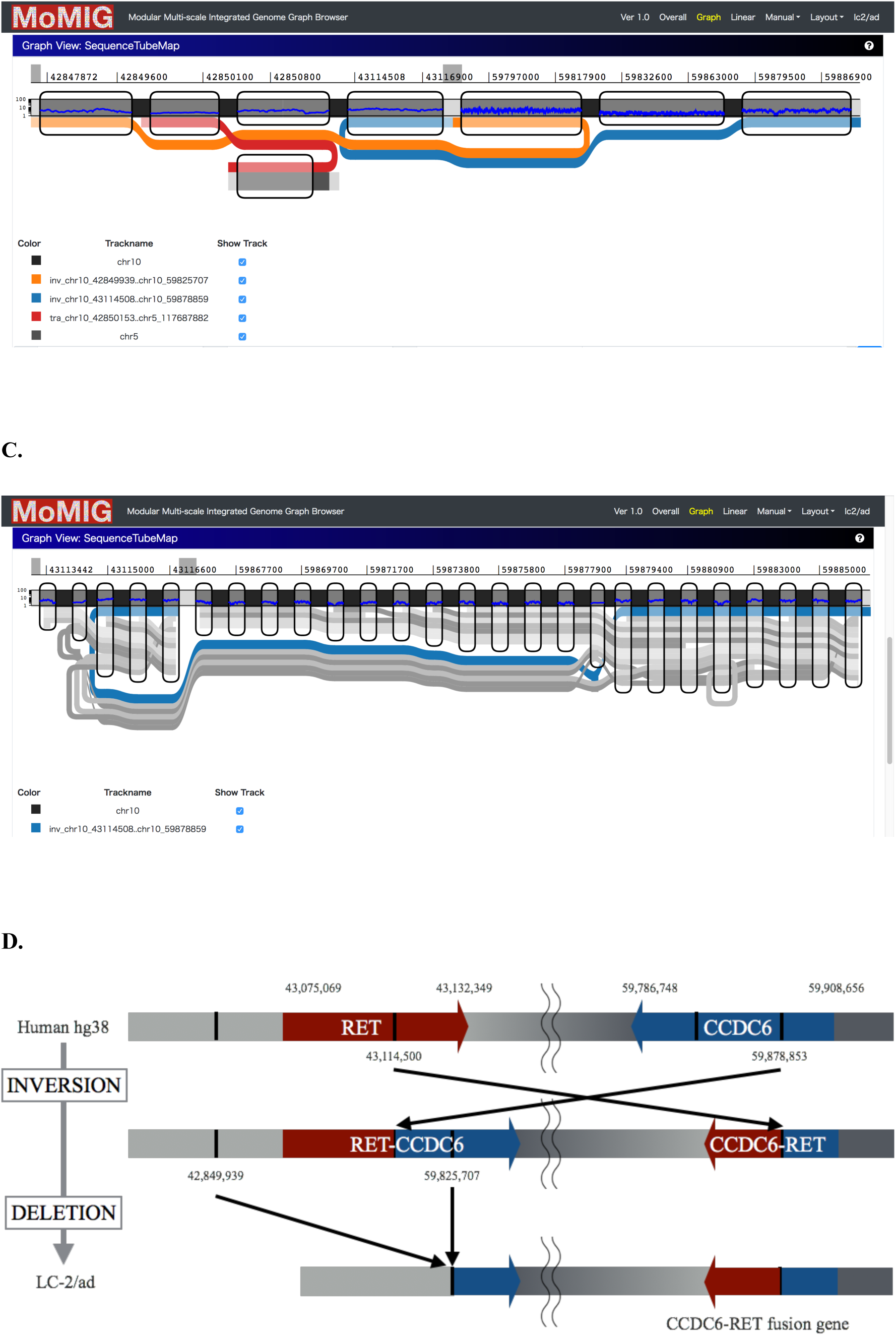
Example of CCDC6-RET. (A) CCDC6-RET shown in MoMI-G (compressed view). The thickest black line is chromosome 10 (reference genome). Note that the two distinct intervals of chromosome 10 are shown, which correspond to the RET (left interval) and CCDC6 (right interval) genes. The blue line represents an inversion identified by Sniffles, showing the CCDC6-RET fusion event. The other lines are gene annotations in hg38; the orange, red, and purple lines indicate two isoforms of RET, and the brown line is CCDC6. (B) CCDC6-RET with read alignments shown as grey lines. Further, some alignments do not support the inversion, suggesting that CCDC6-RET is heterozygous. (C) The entire picture of the inversion that caused CCDC6-RET. This inversion is too large to span by a single read; thus, it was identified as two independent fusion events at both the ends of the inversion, which would be difficult to understand if the two fusion events are visualized separately. The red line is a translocation that was not analyzed in this study. (D) Putative evolution process of LC-2/ad at the CCDC6-RET site. First, a long inversion generated two fusion genes, CCDC6-RET and RET-CCDC6. Second, a large deletion caused the loss of RET-CCDC6.

Further, we found that the large inversion was flanked by two small deletions. These deletions are explained by a single deletion event following the large inversion event (Fig. 2D). The loss of the RET-CCDC6 fusion gene corresponds to the two small deletions on GRCh38. A simple explanation is that a deletion occurred after the inversion event, but not vice versa, in favor of the smaller number of mutation events.

Next, we attempted to estimate the breakpoints of the large inversion before the deletion occurred. There were two possible scenarios for the positions of the two breakends of the large inversion. The first is that RET-CCDC6 and CCDC6-RET were generated by a large inversion and then RET-CCDC6 was lost. The second is that CCDC6 was first broken by a large inversion, and a subsequent small deletion led to CCDC6-RET. Previous studies support the former scenario. First, the RET gene often tends to be disrupted in thyroid cancer by paracentric inversion of the long arm of chromosome 10, or by chromosomal fusion [19]. Second, in a previous study, two clinical samples had both RET-CCDC6 and CCDC6-RET in the genome [20]. Both studies suggested that an inversion disrupted both CCDC6 and RET, and then a small deletion disrupted RET-CCDC6. We could never recognize these two deletions flanking the large inversion without simultaneously observing both the inversion records in VCF.

#### Nested SVs with alignment coverage

Visualizing nested SVs is necessary for evaluating the output of SV callers. However, most existing genome browsers cannot visualize nested SVs as well as the relationships between them. Genome browsers, including IGV [21], collapse SVs into intervals between breakpoints, and thus the topological relationships between nested SVs are not shown. MoMI-G can visualize nested SVs as a variation graph (Fig. 3).

**Figure 3.**
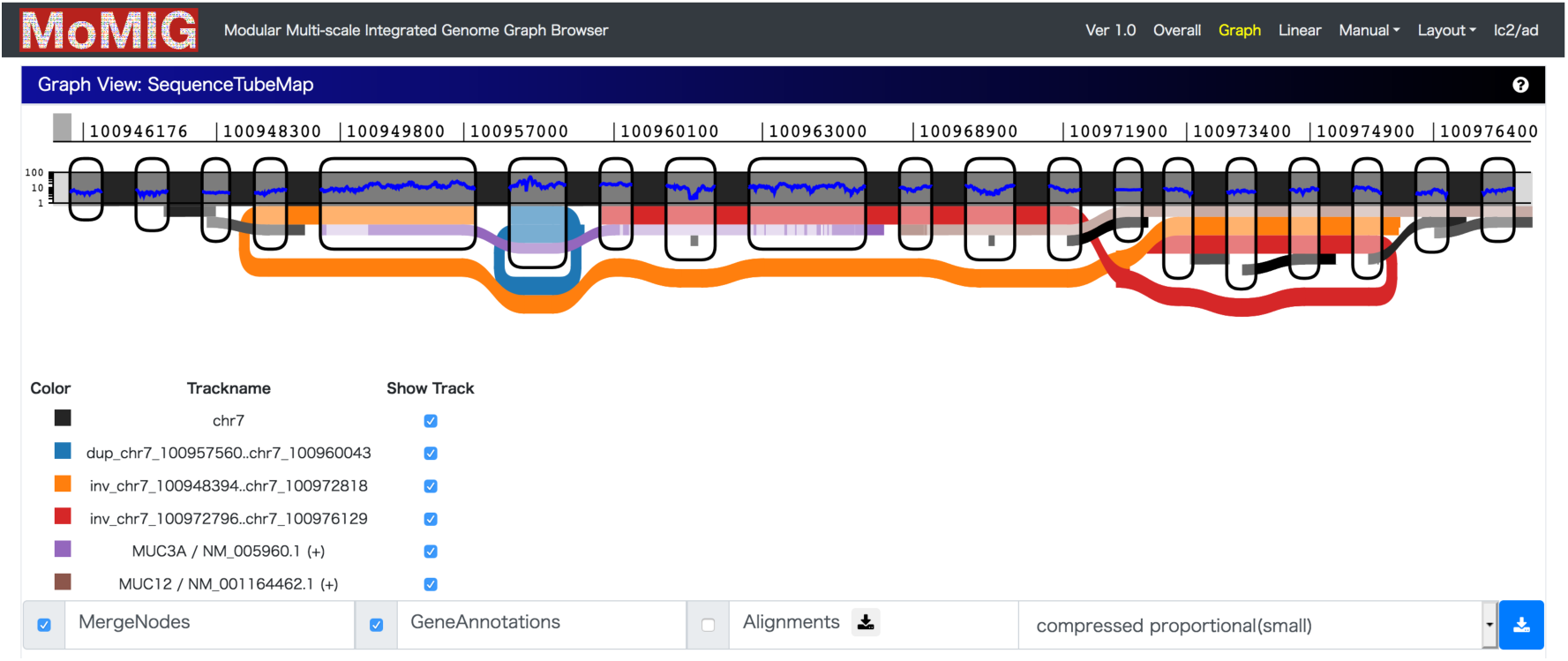
Nested SVs called by Sniffles in LC-2/ad. The thin black lines are repeat annotations. The brown and purple lines are gene annotations. The red and orange lines are an end of an inversion called by Sniffles. There are two possibilities for the genome structure: one is that MUC3A and its flanking region are a duplication, and the internal region of MUC12 is an inverted duplication. The other is that MUC3A and its flanking region are an inverted duplication, and the internal region of MUC12 is a duplication. Several read alignments support the former interpretation. Although SVs called from the Illumina reads did not include any of the SVs shown here, the alignment coverage by the Illumina reads is consistent with both duplications. Note that the y-axis of the blue thin line on the chromosome showing the alignment coverage is logarithmic.

#### Visualizing nested SVs in a pseudodiploid genome

We show an example of nested SVs in a pseudodiploid genome visualized using MoMI-G. We downloaded a CHM1 genome with an SV list previously generated in a whole-genome resequencing study with PacBio sequencers from human hydatidiform [17]. The SV list includes insertions, deletions, and inversions for GRCh37/hg19. We converted the BED file of the SV list of CHM1 to a VCF file, and then filtered out deletions of less than 1,000 bp to focus on medium to large SVs. We found nested SVs for which existing genome browsers do not intuitively show the relationships between them (Fig. 4). This example indicates that four insertions and deletions occur in the large inversion.

**Figure 4.**
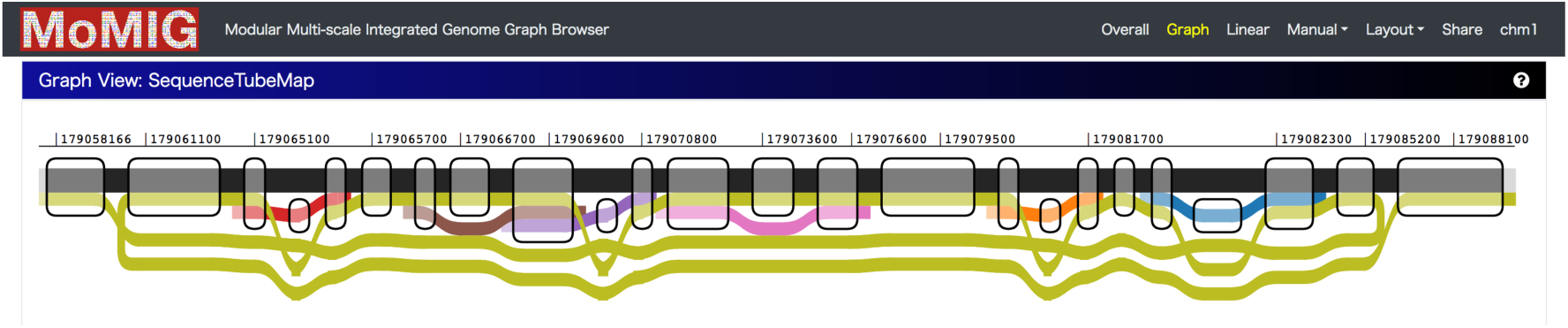
Nested SVs in CHM1. The black line represents a part of chromosome 5, where a large inversion is shown as the brown line. The other lines are smaller SVs included in the large inversion. Because CHM1 is a pseudodiploid genome, all the SVs shown in this figure must be on the same haplotype, although MoMI-G tools assume diploid (polyploid) genomes and show the inner SVs as heterozygous SVs.

### User-interface Design

The optimal way of visualizing SVs might vary. To rapidly explore the distribution of SVs in a genome, users might wish to use Circos-like plots. Other users might intend to focus on local graph structures of SVs that contain a few genes. In another scenario, a user might want to explore individual nucleotides. To address this issue, MoMI-G provides a customizable view in which users can place any combination of view modules. Further, preset view layouts are available for users’ convenience.

#### Enabling easy manual inspection of detected SVs

Manual inspection, which includes determining if an SV is heterozygous or homozygous, confirming what part of a gene is affected by that SV, and determining the reason why an SV is called based on read alignments, is an important part of validating called SVs. As variants called by SV callers increase, the burden of manual inspection also increases, underscoring the importance of visualization both to inspect individual SV calls for filtering out false positives and to ensure that a filtered set of SVs is of high confidence [22,23]. MoMI-G helps with the efficient inspection of SVs by using (1) Feature Table, which is an SV list, (2) Interval Card Deck, which is genomic coordinate stacks, and (3) shortcut keys. The usage is as follows: (1) one can filter SVs using Feature Table, after which SVs are selected, and then (2) the listed variants are stacked on Interval Card Deck at the bottom of the window. In Interval Card Deck, intervals are displayed as cards, and the interval at the top (leftmost) card of the deck is shown on SequenceTubeMap. Each card can be dragged, and the order of cards can be changed. If one double-clicks on a card, the card moves to the top of the deck. A tag can be added for a card for later reference. Further, a card can be locked to avoid unintended modification or disposal, and the gene name can be input with autocompletion for specifying the interval of a card.

When the interval to view is changed, only part of the view that needs an update is re-rendered, whereas most genome browsers working on web interface require rendering the entire view. Interval Card Deck enables the rapid assessment of hundreds of intervals. Moreover, deciding whether an SV should be discarded or held becomes easier with shortcut keys. After all SVs are inspected, a set of SVs held on the Interval Card Deck is obtained, which might be a set of interesting SVs or a set of manually validated SVs. MoMI-G enables the rapid inspection of hundreds of SVs, providing a tool for validating hundreds of SVs or for selecting interesting SVs.

#### Input requirements

MoMI-G inputs an XG format as a variation graph. Users can specify a GAM file with an index that contains read alignment. They can convert a BAM file into GAM using MoMI-G tools or can generate the GAM file on their own. When a BED file of genes is provided, users can specify a genomic interval by gene name. A configuration file is written in YAML. MoMI-G also accepts bigBed and bigWig formats [24] for visualizing annotations (e.g., repeats, genes, alignment depth, and GC content) on the reference genome. The bigBed and bigWig need to be extended for genome graphs in the future. The list of formats that are accepted by MoMI-G is shown in Table 1.

**Table 1.** MoMI-G data files.

| File type | Extension | Description |
| --- | --- | --- |
| A succinct index of variation graphs | .xg | Variation graphs displayed in MoMI-G. |
| Graphical alignment/map | .gam | Read alignment. (optional) |
| Comma-separated values | .csv | SV list for chromosome-scale view. (optional) |
| Browser extensible data | .bed | Used for converting gene names to genomic intervals. Also used for autocompletion of gene names. (optional) |
| Compressed binary indexed BED | .bb | Annotations. (optional) |
| Compressed binary indexed wiggle | .bw | Annotations. (optional) |

## DISCUSSION

We developed a genome graph browser, MoMI-G, that visualizes SVs on a variation graph. Existing visualization tools for SVs show either one SV at a time, or all SVs together; the former does not allow the understanding of the relationships between SVs, whereas the latter is useless when the target genome is very large and the whole variation graphs are too complicated to view in a single screen (i.e., the hairball problem). MoMI-G allows viewing only part of the genome, which resolves the hairball problem, while providing an intuitive view for multiple SVs, including large and nested SVs. Further, MoMI-G enables the manual inspection of complex SVs by providing integrated multiple view modules; users can filter SVs, validate them with read alignments, and interpret them with genomic annotations.

We used vg as a server-side library and SequenceTubeMap as a client-side library for subgraph retrieval and visualization of genome graphs, because to our knowledge, these are the only combinations that are publicly available. We found that significant amounts of engineering efforts are required for an even better user experience. For example, vg is a standalone command line application that exits immediately after the given query is processed; therefore, it does not have a function to keep the succinct index on memory for later use; every time only part of the genome is retrieved, the entire index of several gigabytes is loaded, which is unnecessary overhead. SequenceTubeMap displays inversions and duplications as loops; however, we found that new users occasionally find it difficult to recognize the connections between nodes. Visualizing SVs is still an open problem.

The currently available tools and formats for SV analysis have many problems. First, different SV callers output different VCF records even for the same SV. For example, depending on SV callers, an inversion with both boundaries identified at a base pair level is represented by one of the following: (1) a single inversion record, (2) two inversion records at both ends, (3) two breakend records at both ends, or (4) four breakend records at both ends (a variant of (3), but the records are duplicated for both the strands). Thus, developing a universal tool for variant graph construction is difficult. Second, certain types of nested SVs, such as an insertion within an insertion, are impossible to represent in a VCF file without tricks, although variant graphs can easily handle these SVs. Therefore, generating a variation graph from a VCF file including SVs is not ideal. We need an SV caller that directly outputs variation graphs.

Fostering the ecosystem around variation graphs is important for delivering their benefits to end users, as noted in the ecosystem around the SAM/BAM formats that spurred development of production-ready tools for end users. MoMI-G is the first step toward such a goal, because the availability of tools ranging from upstream analysis, such as read alignment to visualization, is critical for the entire ecosystem.

This is the one million genome era that requires rapid and memory-sufficient data structure to allow alignments and store haplotype information. Because most parts of the genome are shared between individuals, we need to focus on differences for reducing computation time and resources. Therefore, genome graph is considered promising, especially for human variation analysis. Moreover, new visualization methods as well as genome analysis methods are required. Genome graph browsers should be able to handle even thousands of genomes in the near future. MoMI-G is a step forward for visualizing genome graphs and could allow the development of new algorithms on genome graphs.

## METHODS

### Datasets

High-molecular-weight (HMW) gDNA was extracted from lung cancer cell line, LC-2/ad by using a Smart DNA prep(a) kit (Analykjena). WGS data were produced from MinION 1D sequencing (SQK-LSK108), MinION 1D^2 sequencing (SQK-LSK308), and MinION Rapid sequencing (SQK-RAD003). For MinION 1D sequencing, 4 µg HMW gDNA was quantified using Tape Station. DNA repair was performed using NEBNext FFPE DNA Repair Mix (M6630, NEB). End-prep was performed using NEBNext Ultra II End Repair/dA-Tailing Module (E7546L, NEB). Adapter ligation was performed using NEBNext Blunt/TA Ligase Master Mix (M0367L, NEB) and the Ligation Sequencing Kit 1D (SQK-LSK108, Oxford Nanopore Technologies). Libraries were sequenced for 48 h with MinION (R9.5 chemistry, Oxford Nanopore Technologies). For MinION 1D^2 sequencing, the protocol was the same as that for 1D excluding adapter ligation by using Ligation Sequencing Kit 1D^2 (SQK-LSK308, Oxford Nanopore Technologies). The library for MinION Rapid sequencing was prepared according to Sequencing Kit Rapid (SQK-RAD003, Oxford Nanopore Technologies).

### Nanopore data alignment

Nanopore WGS data were aligned against the GRCh38 human reference genome by using NGM-LR with “-x ont” option (version 0.2.6) for calling SVs with Sniffles version 1.0.7 [8]. MinION 1D^2 sequencing can produce a 1D^2 read, which integrates the information of a read and its complementary read into one read. MinION 1D^2 sequencing produces two types of fastq files, 1D and 1D^2; there was some redundancy between 1D and 1D^2 reads. Therefore, redundant 1D reads were removed, and only 1D^2 reads were used. Moreover, some redundant 1D^2 reads were found. These reads were removed from 1D^2 files and used as 1D reads. The percentage of the primary aligned reads was 54.7% (Additional file 1: Supplemental Table 2). This is because we did not filter out reads excluding 1D-fail reads of Rapid sequencing for alignment.

### SV calling

Sniffles (version 1.0.7) with a parameter “-s 5” was used to call SVs. The minimum number of supporting reads was determined such that we could detect the known large deletion of CDKN2A [14], but alignment bias for a reference genome reduces the call rate of insertions compared to that of deletions [17,25]. We detected 11,316 records as a VCF file of SVs, including CCDC6-RET, on the nuclear DNA of LC-2/ad cell line (Additional file 1: Supplemental Table 3). Illumina HiSeq 2000 Paired-end WGS data (DDBJ accession number, DRX015205) aligned against GRCh38 by using bwa (version 7.15) [26] were used for calling SVs with Lumpy (version 0.2.13) [27], delly (version 0.7.7) [28] and manta (version 1.0.3) [29]. The results of these SV callers are listed in Additional file 1: Supplemental Table 5. The four lists of SV candidates detected using Sniffles, Lumpy, delly, and manta were merged using SURVIVOR (version 1.0.0) [30] with “5 sv_lists 1000 1 1 0 0” option for clustering SV candidates (Additional file 1: Supplemental Fig. 3). Further, we filtered the merged candidates based on the following three criteria: (1) Remove SVs of 0 bp to 1 kbp in length to focus on large SVs, (2) Remove all insertions and non-canonical SV types: INVDUP, DEL/INV, and DUP/INS (technically unnecessary, but we aimed at cross-validating SV candidates from Illumina reads with which insertions and non-canonical SV types are hard to call), and (3) Remove SVs overlapping with the intervals of the 10X default blacklist (https://github.com/10XGenomics/supernova/blob/master/tenkit/lib/python/tenkit/sv_data/10X_GRCh38_no_alt_decoy/default_sv_blacklist.bed) for reducing false positives. After filtering, we obtained 1,790 SV records.

### Constructing a variation graph

We constructed a variation graph, including a reference genome (GRCh38) and an individual genome (either LC-2/ad or CHM1). We developed scripts that we call MoMI-G tools. MoMI-G tools follow the procedure in SplitThreader Graph [31].

1. Extract a breakpoint list from a VCF file generated by Sniffles.
2. Construct an initial variation graph with each chromosome of the GRCh38 primary assembly as a single node.
3. Split nodes every 1 Mbp due to the limitation in the implementation of vg used during the development of MoMI-G; otherwise, vg aborted with an error. The latest version of vg does not have this limitation.
4. For each breakpoint in the breakpoint list, split the node of the graph that contains the breakpoint into two nodes at the breakpoint.
5. Create paths that represent the chromosomes in the reference genome.
6. Create paths that represent SVs, each of which corresponds to one record in the input VCF file. Further, add a node to the graph when the type of SV is an insertion.

The representation of SVs varies between SV callers: the same SV can be described in different ways in the current VCF format. Although vg has a “construct” subcommand that constructs a variation graph from a pair of a reference genome and a VCF file, “vg construct” is incompatible with the output of SV callers we know of for the following reason. For example, an INV record in VCF means either a single inversion (both ends included) or only one end of the inversion, depending on the implementation of SV callers. We wrote custom scripts, MoMI-G tools, for Sniffles.

In general, to display read alignments, we recommend aligning against a variation graph by directly using “vg map” instead of converting read alignments against the linear reference genomes into the GAM format. Nevertheless, we intended to inspect the results of Sniffles; therefore, we wrote a custom script to convert alignments against the linear reference (BAM file) into a GAM file by using “vg annotate,” during which CIGAR is lost; “vg annotate” was originally designed for placing annotations in BED/GFF files on paths on variation graphs and not for alignments. One possible solution to this issue is making SV callers graph-aware.

### Inspection of SV candidates

We modified and integrated SequenceTubeMap into MoMI-G so that it can visualize a variation graph converted from SVs that shows the difference between a reference genome and an individual genome. The modifications made to SequenceTubeMap are shown in Additional file 1: Supplemental Table 6. One can click on the download button for downloading an SVG image generated by SequenceTubeMap so that a vector image can be used for publication.

### Visualizing annotations

Ideally, MoMI-G provides annotations on variation graphs. However, annotations available in public databases are for the linear reference genome. MoMI-G can display annotations in the bigWig/bigBed formats. In particular, for human reference genomes, GRCh19/hg37 and GRCh38/hg38, MoMI-G provides an interface for retrieving Ensembl gene annotations from the Ensembl SPARQL endpoint [32] via SPARQList REST API (https://github.com/dbcls/sparqlist). The orientation of genes is shown in the legend of SequenceTubeMap. Further, if one clicks on a gene name, the website of the gene information in TogoGenome [33] opens.

### Miscellaneous modules

Threshold Filter: Threshold Filter has two use cases: First, one can toggle checkboxes to select whether to show inter-chromosomal SVs and/or intra-chromosomal SVs; second, one can filter SVs with a slider based on the custom priority (possibly given by SV callers) of each SV.

Annotation Table: Annotation Table shows all annotations that are displayed on the SequenceTubeMap. Moreover, annotations can be downloaded as a BED file.

Linear Genome Browser: To provide a compatible view of a selected genomic region, we integrated Pileup.js [34] into MoMI-G.

### Backend server

The MoMI-G backend server caches subgraphs once a client requests a genomic interval on a path. It then retrieves annotations from bigWig and bigBed with a range query and provides JSON API with which the client can make queries.

## DECLARATIONS

### Ethics approval and consent to participate

Not applicable.

### Consent for publication

Not applicable.

### Availability of data and materials

Newly obtained long-read sequencing data of LC-2/ad were deposited in the DDBJ with accession numbers DRA007941 (DRX156303-DRX156310). Datasets included in this article are also provided in the database, DBTSS/DBKERO [35]. The source code of MoMI-G is available at https://github.com/MoMI-G/MoMI-G/ under the MIT license. It is written in TypeScript and was tested on Linux and Mac operating systems. Further, we have included an example dataset and annotations as a Docker image.

### Authors’ contributions

All authors were involved in study design; TY wrote all code of MoMI-G; Y. Sakamoto preformed MinION sequencing and SV analysis on LC-2/ad; TY, Y. Sakamoto, and MK drafted the manuscript; MS and Y. Suzuki supervised the sequencing analysis; and MK supervised the study. All authors read and approved the final manuscript.

## Acknowledgements

We thank all testers who provided testing and feedback for MoMI-G. We also thank Kazuyuki Shudo at the Tokyo Institute of Technology and Toshiaki Katayama at the Database Center for Life Science (DBCLS) of Research Organization of Information and Systems (ROIS), Japan, for their useful and insightful discussions, Sarun Sereewattanawoot and Ayako Suzuki at the University of Tokyo for initial contributions to Illumina read alignment.

## ADDITIONAL FILES

Additional file 1: Additional figures and tables referenced in the manuscript. (PDF)

