## Additional file 1 for "MoMI-G: Modular Multi-scale Integrated Genome Graph Browser"

### SUPPLEMENTAL FIGURES

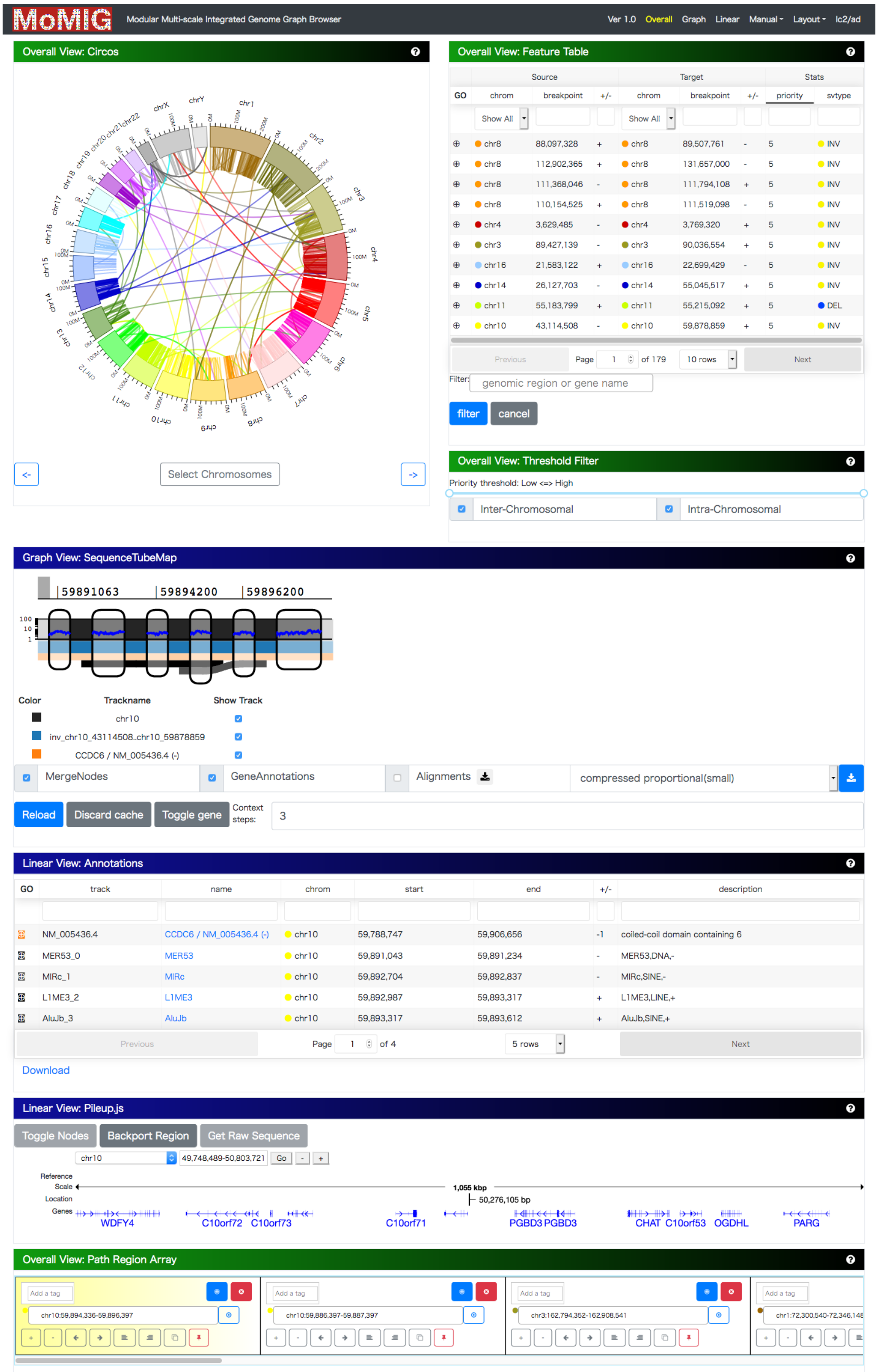

**Supplemental Figure 1. A representative screenshot of MoMI-G with all view modules.**

A

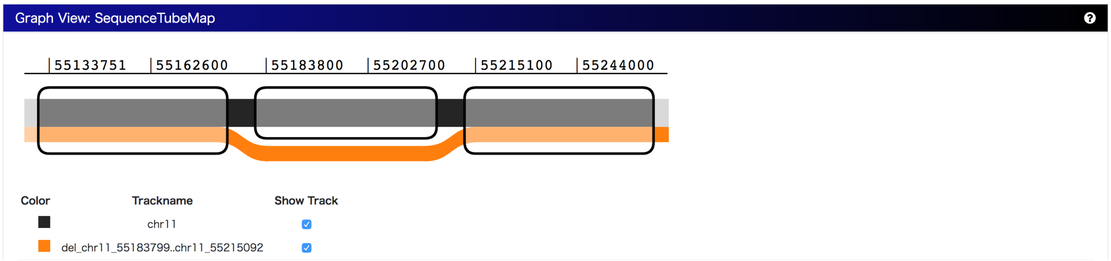


B

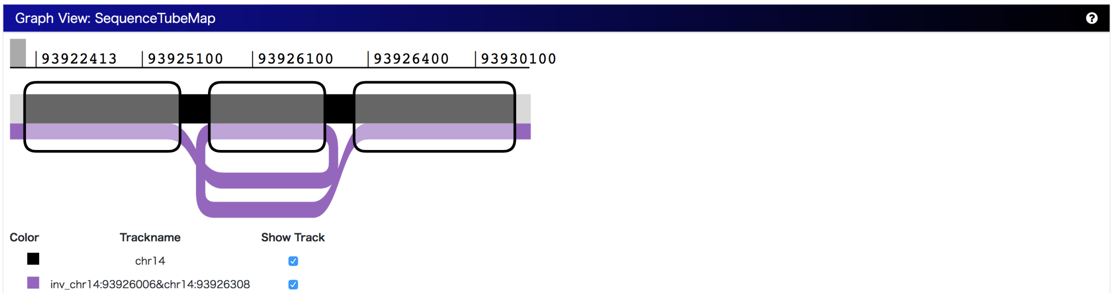


C

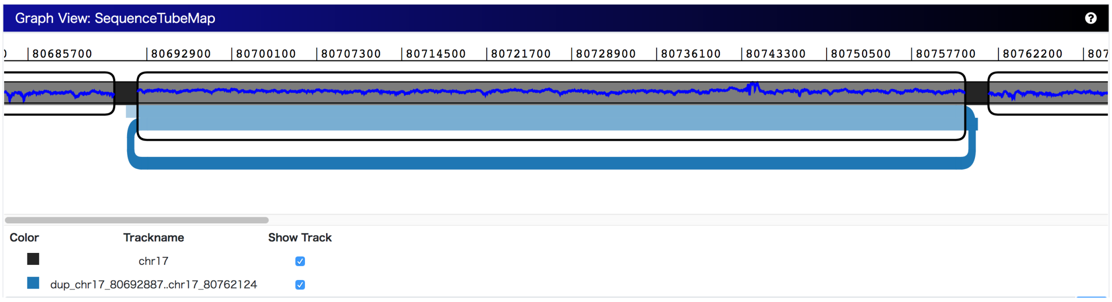


**Supplemental Figure 2. Examples of a deletion, balanced inversion, and duplication**

(A) Deletion: The orange line indicating a deletion starts before one breakend of the deletion, passes through the middle node that indicates the deleted sequence, and then proceeds for the sequence flanking the deletion.

(B) Balanced Inversion: The purple line indicating a balanced inversion includes flanking sequences on both breakpoints of the inversion.

(C) Duplication: The blue line indicating a duplication passes through the node twice, suggesting that the sequence of the node is duplicated. The line in the node might terminate if the node is interrupted by other SVs.

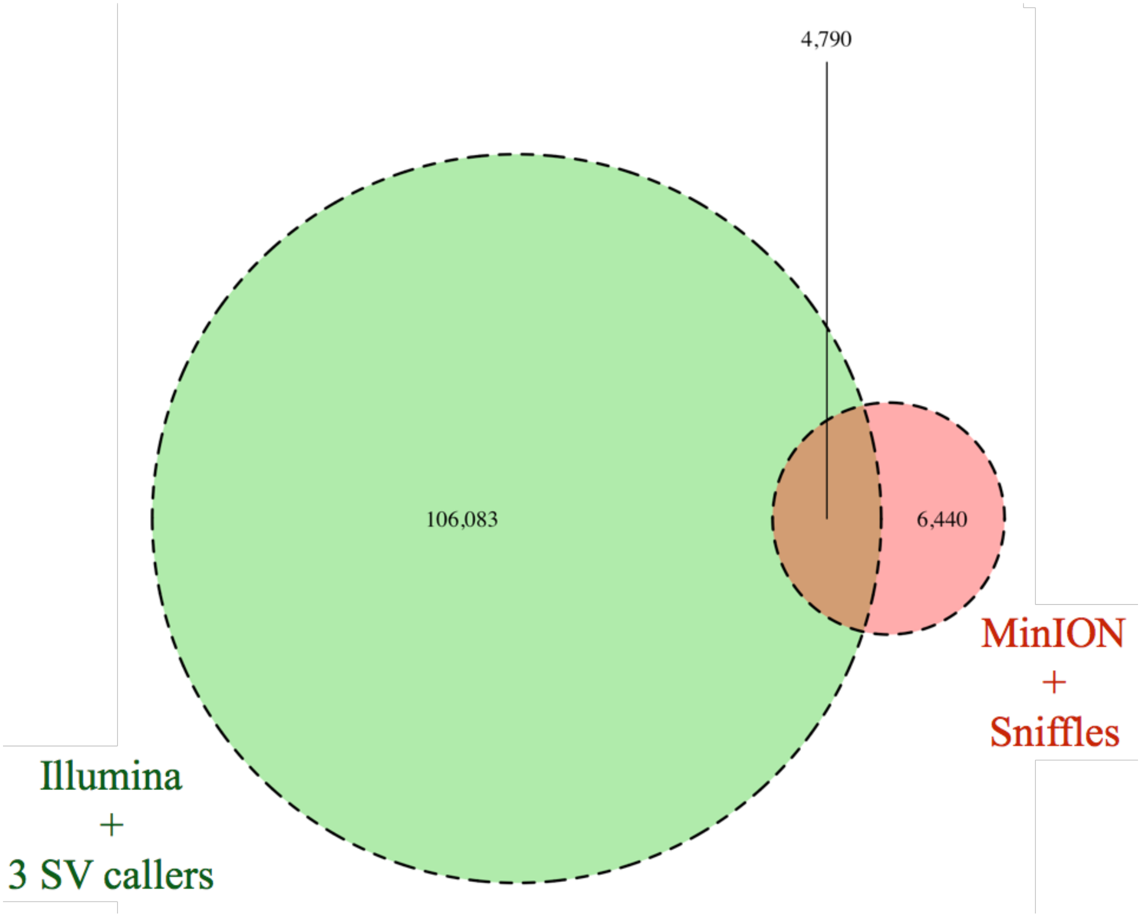


**Supplemental Figure 3. Merged SVs from four SV callers by using SURVIVOR**

The green circle describes the count of SVs called by Illumina, whereas the red circle presents the count called by MinION.

### SUPPLEMENTAL TABLES

#### Supplemental Table 1. List of MoMI-G features

| Module group | View module | Description |
|---|---|---|
| Chromosome-scale View | Circos Plot | Observe the distribution of SVs |
| | Feature Table | Select and filter SVs |
| | Threshold Filter | Filter SVs |
| | Interval Card Deck | Accumulate genomic intervals for inspection |
| Gene-scale View | SequenceTubeMap | Visualize a subgraph of a variation graph |
| | Interval Card Deck | Switch between the intervals |
| Nucleotide-scale View | Linear Browser | Visualize a linear genome with annotations |
| | Annotation Table | Show annotations |
| | SequenceTubeMap | Visualize nucleotides of a variation graph |
| | Interval Card Deck | Switch between the intervals |

#### Supplemental Table 2. Summary of the Oxford Nanopore sequencing data of LC-2/ad

| | |
|---|---|
| Number of reads | 3,310,982 |
| Coverage | 12.8× |
| Percentage of aligned reads | 57.9% |
| Average length of aligned reads | 16,032.2 bp |
| Average identity of aligned reads | 83.0% |

**Supplemental Table 3. Called SVs in LC-2/ad nuclear DNA from Nanopore reads**

| SV Types | Called SVs |
|---|---|
| DEL | 6,366 |
| DUP | 323 |
| INS | 4,449 |
| INV | 99 |
| INVDUP | 1 |
| TRA | 79 |
| DEL/INV | 12 |
| DUP/INS | 3 |

**Supplemental Table 4. Breakends in the opposite end of the CCDC6-RET inversion**

| Sequencer, aligner, and SV caller | Start | Stop |
|---|---|---|
| ONT MinION, NGM-LR + Sniffles | chr10: 42,849,939 | chr10: 59,825,707 |
| Illumina HiSeq 2000, BWA + manta | chr10: 42,849,938 | chr10: 59,825,707 |
| Illumina HiSeq 2000, BWA + delly | chr10:42,849,938 | chr10:59,825,707 |
| Illumina HiSeq 2000, BWA + lumpy | Not called | Not called |

**Supplemental Table 5. Called SVs in LC-2/ad nuclear DNA from Illumina reads**

| SV types | manta | delly | lumpy |
|---|---|---|---|
| BND | 3,926 | 9,787 | 4,214 |
| DEL | 43,886 | 33,429 | 2,051 |
| DUP | 768 | 3,213 | 1,197 |
| INS | 29,216 | 2,212 | - |
| INV | 576 | 2,055 | 62 |

**Supplemental Table 6. The modifications made to the original SequenceTubeMap**

| Original implementation | Modified implementation |
|---|---|
| Paths are displayed in the same order as they appear in the input. | A selected chromosome path is by default drawn as a straight horizontal line regardless of the path order in the input. |
| It was impossible to display genes and paths in different colors. | Genes, annotations, chromosomes, and variants can have different colors for easily distinguishing them. |
| The constant for the linear node width mode was too large for visualizing large SVs. | We implemented another mode suitable for visualizing large SVs. |
| No bigWig/bigBed support. | We added bigWig/bigBed support. |
| Non-consecutive genomic intervals are difficult to recognize. | We inserted a shaded gap between distant genomic intervals. |
